# Fixation of a homozygote and dynamical characterization of the Price equation

**DOI:** 10.64898/2026.04.22.720281

**Authors:** József Garay, Tamás F. Móri

**Author notes:** Corresponding author: József Garay, Konkoly-Thege M. út 29-33, H-1121 Budapest, Hungary. **Author Contributions:** Conceptualization: József Garay, Tamás F. Móri; Methodology: József Garay, Tamás F. Móri; Formal analysis and investigation: Tamás F. Móri; Writing - original draft preparation: József Garay, Tamás F. Móri; Writing - review and editing: József Garay, Tamás F. Móri; Supervision: József Garay.

## Abstract

Price equation and genotype dynamics are two methods for studying the fixation of one allele by natural selection in a diploid population. There are two strict monotonicity conditions that imply the fixation of one allele. The genotype dynamics is called Haldane monotone if the relative frequency of *one allele* strictly increases along all solutions of the genotype dynamics, so this allele is fixed. In this paper, we show that the genotype dynamics is Haldane monotone if and only if the right-hand side of the Price equation is always strictly positive. The other strict monotonicity condition requires that the relative frequency of *a homozygote* strictly increase according to the genotype dynamics. For example, in a model where the genotype dynamics is governed by interactions between individuals, *the cost-accepting homozygote is fixed by natural selection if the other genotypes always receive a smaller average gain from all interactions than the cost-accepting homozygote*. Both monotonicity conditions require that the interaction is not well-mixed in the population. These two conditions are not equivalent.

In addition, we give a non-monotonicity condition, which also implies the fixation of a homozygote. The fixation of a homozygote depends on the phenotypic payoff of the interaction, the genotype-phenotype mapping, and the interaction scheme. In a sexual population, the interaction scheme of siblings depends on the mating system, and so do the conditions of fixation of the cost-accepting homozygote. We present examples showing that if we only change the monogamous mating system, assuming panmixing or mating assortativity, then the condition for the fixation of the cooperator homozygote is *b* > 2*c* and *b* > *c*, respectively.

## 1. Introduction

One of the puzzles of evolutionary biology is why one individual helps another at the expense of itself. Altruism is a one-person game, where the focal individual either helps her companion in need at her own expense or does not help (Hamilton 1970). Here only the actor has strategy. The prisoners’ dilemma is a two-person game, and when the cooperator interacts with the defector, the cooperator pays the costs while the defector only benefits from it (Axelrod & Hamilton 1981). In both cases, the cost-avoidant (e.g. selfish or defector) individual has a relative advantage over the cost-accepting (e.g. altruistic or cooperator) individual who pays the cost, which does not seem reasonable from an evolutionary perspective.

In this paper, we only focus on one of the possible explanations of the evolutionary advantage of the cost-accepting behavior, namely, the kin selection. In the kin selection theory (p.44 in Haldane 1955, Hamilton 1964a,b, Maynard Smith 1964) the direct benefit is replaced by indirect one. The evolutionary success of the altruistic gene is increased whenever an individual having the altruistic gene promotes the survival of its sibling who also carries the same altruistic gene, thus the altruistic individual indirectly increases the frequency of its altruistic gene. The well-known mathematical formulation of this indirect effect is the classical Hamilton’s rule (Hamilton 1964): the altruistic allele will spread if *rb* > *c*, where *b* is the fitness benefit to the diploid recipient, *c* is the fitness cost to the diploid actor and *r* is the coefficient of genetic relatedness between the interacting individuals. Hamilton’s rule claims that under this condition the altruistic allele is globally fixed by natural selection.

There are a couple of methods to handle the fixation of an allele in kin selection theory (for a review see van Veelen et al. 2017). Here we only focus on the relationship between the following two deterministic methods and only provide background information to the extent absolutely necessary.

### Method 1. The Price equation

Price (1970) originally described the frequency change of a given allele in two consecutive diploid generations. The selection situation is as general as possible in the sense that we only need to know the number of each genotype in the next generation. In other words, the Price equation is valid regardless of how we specify the number of individuals of each genotype in the next generation. The Price equation has played a central role in kin selection theory (Frank 1995). The following observations from the Price equation literature are relevant to this article. Although the Price equation is very general, using the Price equation, Queller (1992) showed that when the costs and the benefits are expressed as their average effects on genes, Hamilton’s rule holds. Recently, van Veelen (2025) derived a generalized Price equation. He also considered individual fitness dependent on allele frequency. Moreover, van Veelen (2025) pointed out that the form of the generalized Hamiltonian law depends on the fitness definition. Here we follow Price, and make no assumptions about the number of individuals of each genotype in the next generation. Furthermore, there is a controversial opinion, that a disadvantage of the Price equation is its dynamical insufficiency (e.g. Frank 1995, van Veelen et al. 2012). Here we will examine the Price equation from the perspective of genotype dynamics.

### Method 2. Genotype dynamics

According to Maynard Smith (1964), kin selection occurs when the closest relatives interact. The closest relatives, i.e., full siblings, are found in monogamous families. Haldane (1924) introduced “familial selection” 100 years ago, where the struggle of life takes place between full siblings for food. We developed a population genetic model to investigate familial selection in a sexual population (Garay et al. 2019), where a homozygote has either a cost-accepting or cost-avoidant phenotype. To investigate our familial selection model, we developed the following population genetics methodology. We adapted the verbal definition of classical ESS (Maynard Smith & Price 1973) to the diploid, sexual, Mendelian population (Garay et al. 2019, 2025). We used genotype dynamics (Garay et al. 2019, 2025 Haldane & Jayakar 1965, Hull 1964) to investigate the fixation of a homozygote. We also introduced the concept of Haldane’s monotonicity, where the frequency of an allele strictly increases along all solutions of the genotype dynamics (Garay et al. 2025), and consequently the homozygote of this allele is fixed. The quality of any method depends on what it can be used for and whether or not it is able to provide new results or insight. We have already examined selection situations where altruistic help (Garay et al. 2019, 2023, 2024), and where prisoner’s dilemma (Garay et al. 2025) takes place between full siblings, and the pure strategies are determined by two alleles at an autosomal locus. One of the novelties of our population genetics model is that the fixation conditions of the altruistic or cooperative allele depend on the genotype-phenotype mapping (Garay et al. 2019, 2023, 2024, 2025), and the mating system (Garay et al. 2023). Furthermore, using genotype dynamics, we established that in the prisoner’s dilemma, homozygous fixation of the cooperator and homozygous fixation of the defector, as well as coexistence of the cooperator and defector siblings, are possible. To our knowledge, no previous results similar to ours have been published. Based on the above results, we can state that population genetic methodology can provide new insights into kin selection.

There are studies in the literature where the conditions for the fixation of a given allele (or homozygote) are different in different selection situations (e.g. Garay et al. 2023, 2025, van Veelen 2025, Allen et al. 2024). From a mathematical point of view, this makes perfect sense. In recent decades, however, Hamilton’s rule has seemed to be universally valid. The claim to universality has raised two problems. Any universal rule is independent of the method of derivation. Thus, our first questions arises: Can the two methods listed above give the same sufficient condition for single-allele fixation? If the answer is yes, then the next question arises: Which method is more widely applicable? In other words, can genotype dynamics give any general condition for single-allele fixation that Price’s equation cannot?

To answer the above two questions, we first recall the details of our population genetic method and give a new general condition for the fixation of cost-accepting behavior. We then focus on the relationship between the Price model and population genetic methods. We then present an illustrative example. Finally, we discuss the results.

## 2. Population genetics method

For the convenience of the reader, we recall the details of our methodology used here (Garay et al. 2019, 2023, 2024, 2025). We emphasize that the tools of our methodology are general in the sense that the proofs of the results cited here are not based on the details of Haldane’s familial selection (see proofs in Garay et al. 2019, 2025). Furthermore, we conclude this section with two new general conditions for the fixation of a single homozygote.

Let us consider a sufficiently large diploid population of *N* individuals. The females and males differ only in sex, in a 50/50 ratio, and no sexual selection. We follow the change of the genotype frequencies over time. We consider two alleles ([a] and [A]) at a Mendelian autosomal locus, so we have three genotypes. Thus the distribution of genotypes belongs to the two-dimensional simplex *S*_3_ = {*x* = (*x*_1_, *x*_2_, *x*_3_) ∈ ℝ^3^: 0 ≤ *x*_*i*_ ≤ 1, *i* = 1, 2, 3, and Σ*x*_*i*_ = 1}. Let *x*_1_, *x*_2_ and *x*_3_ denote the relative frequencies of the genotypes [aa], [aA] and [AA] in the parent population, respectively. Define the *production* functions *z*_1_, *z*_2_, *z*_3_ as the numbers of genotypes [aa], [aA] and [AA] in the next parent population; (*z*_1_, *z*_2_, *z*_3_): *S*_3_ → ℝ^3^. We can assume that they are smooth. We use the term “production”, because in a diploid sexual population, genotypes can have offspring with different genotypes, e.g. different homozygous parents can only have heterozygous offspring. Since the smooth production functions *z*_*i*_(*x*) can be arbitrary, this selection situation is quite general. Note that Price (1970) already considered this selection situation (see the next section).

We adapt the verbal definition of the classical ESS (Maynard Smith & Price 1973) to this selection situation. We say that a homozygote [aa] is evolutionary stable (Garay et al. 2019), or formally, *x*^*^ = (1,0,0) is an *evolutionarily stable genotype distribution* (ESGD) if *z*_1_(*x*) > *x*_1_ Σ_*j*_ *z*_*j*_ (*x*) for all *x* sufficiently close to *x*^*^. The concept of ESGD is based on the fundamental principle of Darwinism: the relative frequency of a given genotype increases if its production exceeds its proportional share of the total production of the entire population; in other words, if its relative advantage over the entire population is positive (Alós-Ferrer & Hofbauer 2022). To convince the reader that the ESDG concept is not a mere formalism, we recall the concept of Haldane’s arithmetic (Garay et al. 2023), which states that the relative frequency of an altruistic gene increases if, on average, it has ‘saved’ more than one copy of itself in its lineage by ‘self-sacrifice’.” We found that Haldane’s arithmetic is equivalent to the definition of ESGD. The main difference between Hamilton’s rule and Haldane’s rule is that the latter can guarantee the fixation of the altruistic allele through homozygous fixation when the interaction between siblings is given by a nonlinear game (Garay et al. 2023), while Hamilton’s rule cannot.

Analogously to the replicator dynamics, the change in the diploid state of the whole population is expressed as the difference between the production of the given genotype and the total production of the population, more precisely

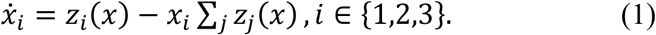

This dynamics is the so called *genotype dynamics* (for a detailed derivation, see SI C in Garay et al. 2019, see also Garay et al 2025). The genotype dynamics follows the change of genotype distribution from generation to generation. Note that in a sexual population, if neither genotype is lethal then the pure heterozygous [aA] state (0,1,0) is not a rest point of the genotype dynamics, and furthermore, none of the edges of the simplex is invariant since the Mendelian genetic system generates all genotypes when both alleles are present.

We proved that if the vertex *x*^*^ = (1,0,0) is an ESGD, then it is a locally asymptotically stable equilibrium point of the genotype dynamics (1) (see SI C in Garay et al. 2019). Note that this theorem allows us to prove the local stability of genotype dynamics using the inequality in the definition of ESGD.

For the global fixation of a homozygote according to the genotype dynamics, we introduced the concept of *Haldane monotonicity* (Garay et al. 2025). We say that the genotype dynamics is *Haldane monotone* if the frequency of the allele [a] in the entire population is a global Lyapunov function of the genotype dynamics (formally 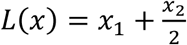 where x ∈ *S*_3_). This means that along all solutions of the genotype dynamics, the relative frequency of the [a] allele always strictly increases. We found that the genotype dynamics is Haldane monotone if and only if 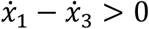 for all *x* ∈ int *S*_3_, that is, in more detail,

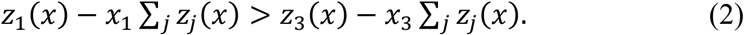

The biological interpretation of this inequality is as follows: The genotype dynamics is Haldane monotone if the relative advantage of the homozygote [aa] over the entire population is greater than that of the homozygote [AA]. It should be noted that global Haldane monotonicity is a rather strong sufficient condition. For example, we found an example where the classical Hamilton’s rule implies global stability of cooperator homozygotes [aa], but the genotype dynamics is not Haldane monotone. For the convenience of the reader, we recall a numerical example of this (see Fig. 1). Example 1 concerns family selection.

**Figure 1.**
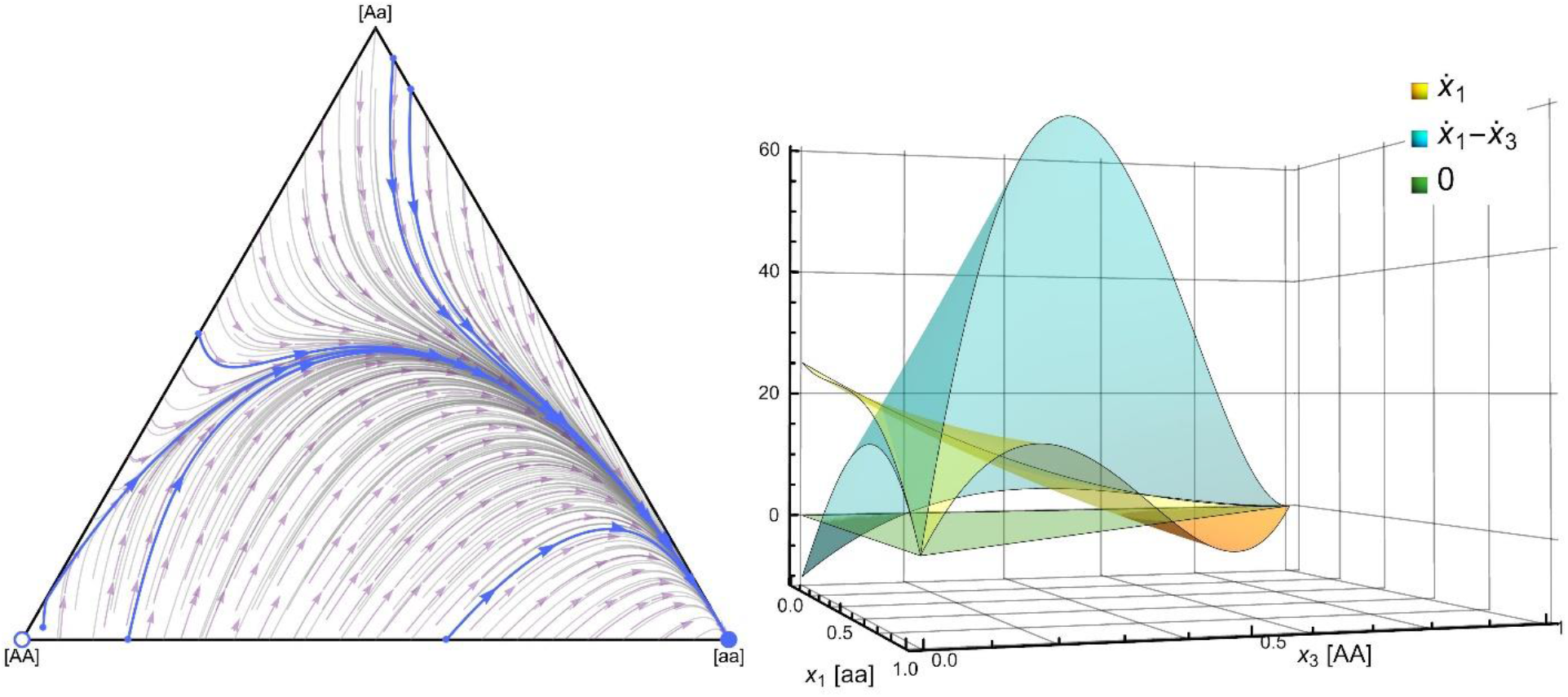
We visualize the genotype dynamics in a panmictic population where the survival probability of juveniles is determined by a survival donation game. Each family has *n* = 8 newborns. In Figure 1A, we consider the case where the cooperator allele [a] is recessive, the basic survival rate *α* = 0.2, the benefit *b* = 0.6 and the cost *c* = 0.1, so the payoff matrix is 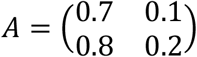. We know that if *b* > 2*c*, then the cooperator homozygote [aa] is ESGD (Garay et al. 2025). Furthermore, the defector homozygote [AA] is a repellor, and this dynamic has no internal rest point, so according to the Poincaré–Bendixon theorem, in this selection situation the classical Hamilton rule implies the global stability of the cooperator homozygote [aa].

**Example 1.** *Panmixia and survival matrix game between full siblings*

As assumed above, we consider a very large diploid population of *N* individuals. Females and males differ only in sex, in a 50/50 ratio between the sexes, and there is no sexual selection. Consider a familal selection situation where siblings play a survival game. The phenotype of each genotype is determined by a Mendelian autosomal locus. The [a] allele is either recessive or dominant to the [A] allele. Let the homozygous [aa] be a cooperator, while the genotypes [aA] and [AA] are defectors. The phenotypic composition of each family is determined by Mendelian inheritance. The sexual population is panmictic. The mating table (Table 1) shows the interaction scheme: *p*_*k*(*ij*)_ denotes the probability that parents with genotypes *i* and *j* will produce offspring with genotype *k*.

**Table 1.**
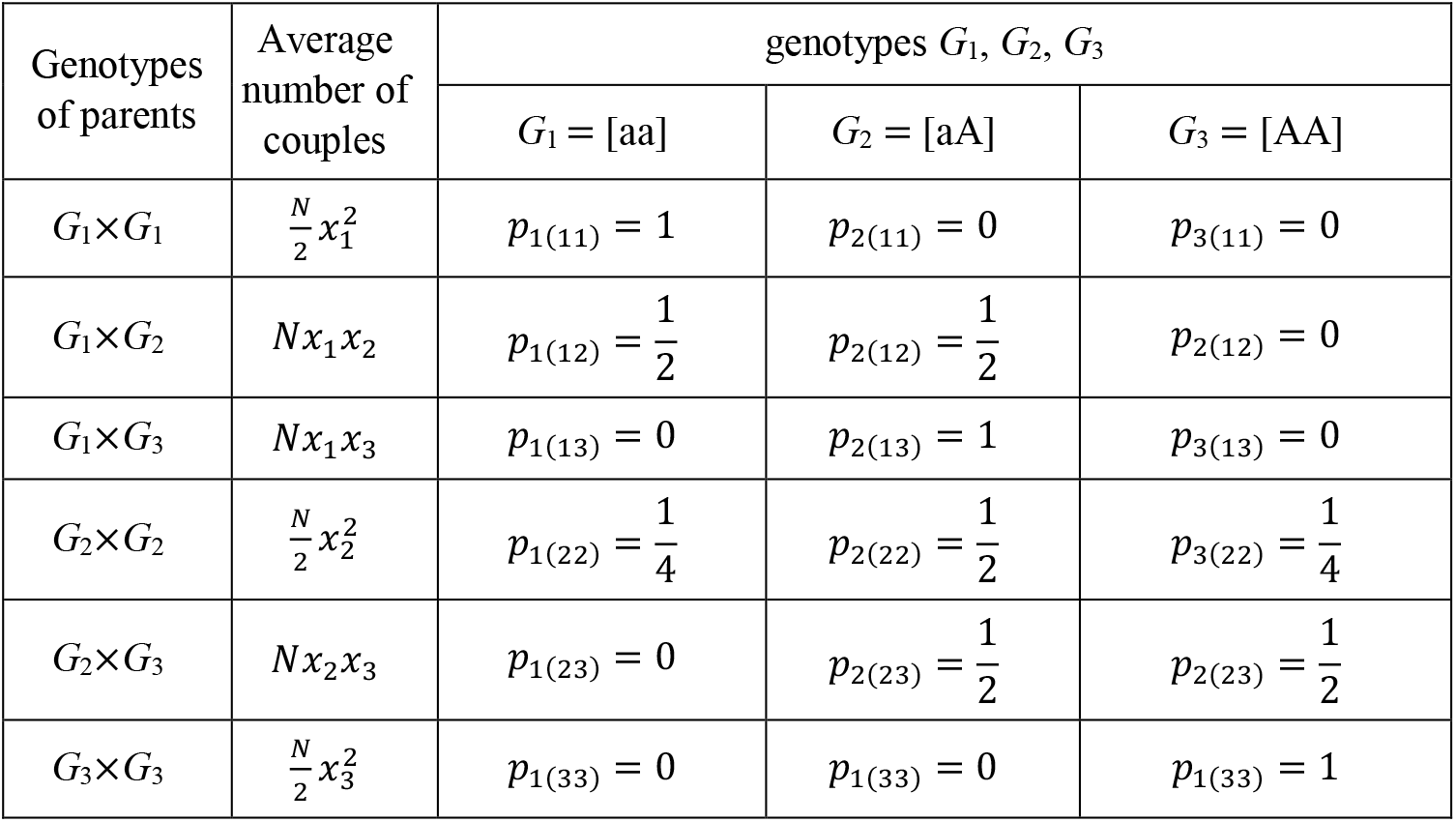
Mendelian inheritance determines the phenotypic composition of individual families, as well as the interaction scheme.

Suppose that in each family there are n newborns, i.e. all females have the same basic fecundity. There are two options that still lead to the same formulation in the long run. Either each offspring plays with all its siblings, or they are randomly paired (suppose n is even) and play so that each only plays only once. The only difference is that in the former model there is an (*n* − 1) multiplier that is not present in the latter one.

Note that the probability of interaction between two sibling genotypes depends on the genotype distribution of the parental population. For instance, the average number of the interactions between two homozygotes [aa] is 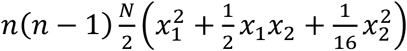, the number of interactions between two different homozygotes is 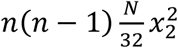 and the number of the interactions between two heterozygotes is 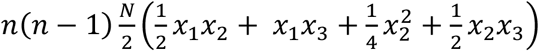, etc. The main point is that the interaction is not well mixed.

The full siblings play a donation survival two person matrix game (each with everyone else), with payoff matrix 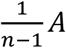, where

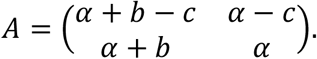

Here *α* is the basic survival rate, *b* is the benefit and *c* is the cost; *α, b, c* are all positive, and the entries of *A* fall between 0 and 1 so that the survival probability remains in the interval [0, 1] even after *n* − 1 interactions. The composition of the different families is random, but as far as we are interested in averages (i.e., expectations), the *n* − 1 interactions played by a sibling can be equivalently substituted by a single one with payoff matrix *A*. The payoff of each individual depends on its genotype and the genotype of its siblings (see Table 2). However, the phenotypic composition of each family depends on the genotypes of the parents. For instance, in a *G*_2_×*G*_2_ family the genotypes *G*_1,_ *G*_2_ and *G*_3_, are born with probabilities of 1/4, 1/2 and 1/4, respectively. As a result, a *G*_1_ juvenile interacts with genotype *G*_1,_ *G*_2_ and *G*_3_ with probabilities 1/4, 1/2 and 1/4, respectively. Denote the strategy of genotype *G*_1,_ *G*_2_ and *G*_3_ by *s*_1_, *s*_2_ and *s*_3_ respectively. Then the survival rate of a focal *G*_1_ juvenile is 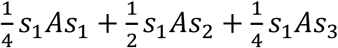. Since the birth probability of a *G*_1_ juvenile in a *G*_2_×*G*_2_ family is 1/4, the average (expected) number of surviving *G*_1_ juveniles in *G*_2_×*G*_2_ families is 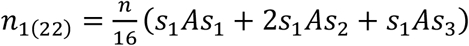. Note that this formula is valid both in the case where each sibling plays with all siblings (with payoff matrix 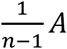) and in the case where each of them plays only one game (with payoff matrix *A*).

**Table 2.** Genotype survival table based on the mating table, for a general survival matrix game. This table corresponds to panmixia, since the average number of different families is determined by the random mating of different parent genotypes. According to Mendelian inheritance, the genotype of the parents determine the phenotypic composition of their family. The strategy of each juvenile is determined by two alleles. If allele [a] is recessive, then genotypes [aA] and [AA] adopt the defector strategy. If allele [a] is dominant, then the genotypes [aa] and [aA] adopt the cooperator strategy.

| Genotypes of parents | Average number of couples | Average number of surviving family members, for genotypes $G_1, G_2, G_3$ | | |
| --- | --- | --- | --- | --- |
| | | $G_1 = [aa]$<br>$s_1$ | $G_2 = [aA]$<br>$s_2$ | $G_3 = [AA]$<br>$s_3$ |
| $G_1 \times G_1$ | $\frac{N}{2} x_1^2$ | $n_{1(11)} = ns_1As_1 = n$ | $n_{2(11)} = 0$ | $n_{3(11)} = 0$ |
| $G_1 \times G_2$ | $Nx_1x_2$ | $n_{1(12)} = \frac{n}{4}(s_1As_1 + s_1As_2)$ | $n_{2(12)} = \frac{n}{4}(s_2As_1 + s_2As_2)$ | $n_{3(12)} = 0$ |
| $G_1 \times G_3$ | $Nx_1x_3$ | $n_{1(13)} = 0$ | $n_{2(13)} = ns_2As_2 = n$ | $n_{3(13)} = 0$ |
| $G_2 \times G_2$ | $\frac{N}{2} x_2^2$ | $n_{1(22)} = \frac{n}{16}(s_1As_1 + 2s_1As_2 + s_1As_3)$ | $n_{2(22)} = \frac{n}{8}(s_2As_1 + 2s_2As_2 + s_2As_3)$ | $n_{3(22)} = \frac{n}{16}(s_3As_1 + 2s_3As_2 + s_3As_3)$ |
| $G_2 \times G_3$ | $Nx_2x_3$ | $n_{1(23)} = 0$ | $n_{2(23)} = \frac{n}{4}(s_2As_2 + s_2As_3)$ | $n_{3(23)} = \frac{n}{4}(s_3As_2 + s_3As_3)$ |
| $G_3 \times G_3$ | $\frac{N}{2} x_3^2$ | $n_{1(33)} = 0$ | $n_{2(33)} = 0$ | $n_{3(33)} = ns_2As_2$ |

Now we can calculate the total number of survivors in the next generation.

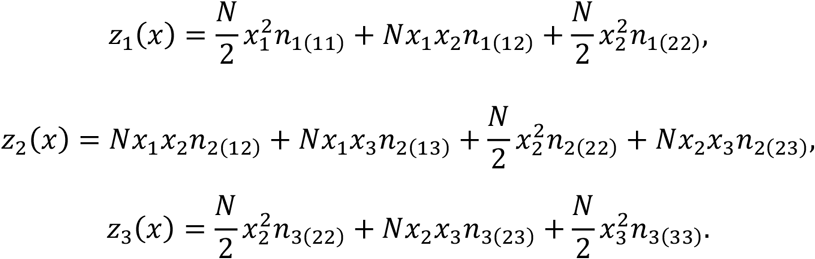

We previously found (Garay et al. 2025) that in this selection situation, the classical Hamilton rule implies the global fixation of homozygote [aa], but the genotype dynamics is not Haldane monotone (see Fig. 1).

In Figure 1B we visualize both monotonicity conditions. This system is not globally Haldane monotone since 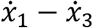 is negative in the neighborhood of the heterozygote vertex (0,1,0). Note that since Price monotonicity and Haldane monotonicity are equivalent, the Price equation cannot show the global stability of homozygote [aa]. Furthermore, our monotonicity condition is only not globally satisfied, since 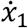 is negative in the neighborhood of the homozygote vertex (0,0,1). However, in an appropriate neighborhood of the pure recessive homozygote [aa] vertex (1,0,0), the system is locally Haldane monotone and our monotonicity condition also holds.

We have now completed the introduction of our population genetic methods which we will need later. Thus, we can provide new conditions for the fixation of an allele by genotype dynamics.

We now formulate general, non-monotonicity conditions that imply global stability of homozygote [aa].

### Theorem 1.

Assume that

a. the homozygote [aa] vertex is attractor (e.g. it is ESGD),
b. the homozygote [AA] vertex is repellor (thus it is not ESGD),
c. apart from the homozygous vertices (1,0,0) and (0,0,1), the genotype dynamics (1) has no rest point.

Then the Poincaré–Bendixson theorem implies the global fixation of the homozygote [aa], i.e. (1,0,0) is a globally asymptotically stable rest point of the genotype dynamics (see SI A1 for the proof).

Note that conditions (a) and (b) are local, and thus easier to verify analytically, while condition (c) is a global condition and thus more difficult to verify analytically. However, in any specific selection situation, each condition can be verified numerically. It is clear that conditions (a–c) are not non-monotonic, since in the interior of the simplex the relative frequency of the homozygote [aa] can decrease.

Furthermore, if both homozygote vertices are repellors, then all genotypes coexist (i.e., there is either at least one asymptotically stable fixed point, or there is an asymptotically stable cycle). If there is no cycle and only one interior rest point exists, then it is globally stable.

Hamilton’s rule and Haldane’s arithmetic are monotonicity conditions. Here we also give a monotonicity condition that guaranties the fixation of a homozygote.

### Theorem 2

(A monotonicity condition on homozygote) Suppose that the relative frequency of the cost-accepting homozygote [aa] always increases, formally

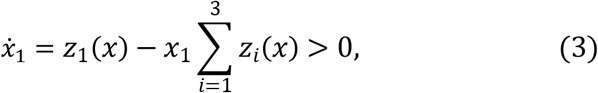

for all *x* ∈ *S*_3_\{(1,0,0), (0,0,1)}. Then the cost-accepting homozygote [aa] is fixed by natural selection (see SI A2 for the proof.)

### Remark 1

The Haldane monotonicity and our monotonicity condition (3) are not the same. For example, in some region of the simplex it may happen that

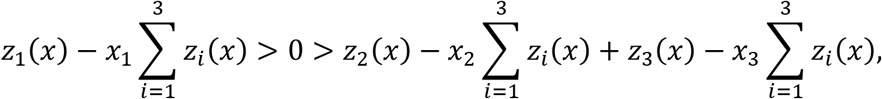

but at the same time

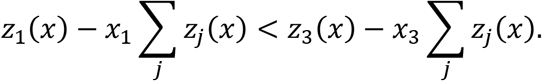

In other words, the relative frequency of homozygote [aa] may increase while that of the allele [a] decreases. Based on Figures 1 and 2, it seems that our examples are also like this and are not globally Haldane monotone.

**Figure 2.**
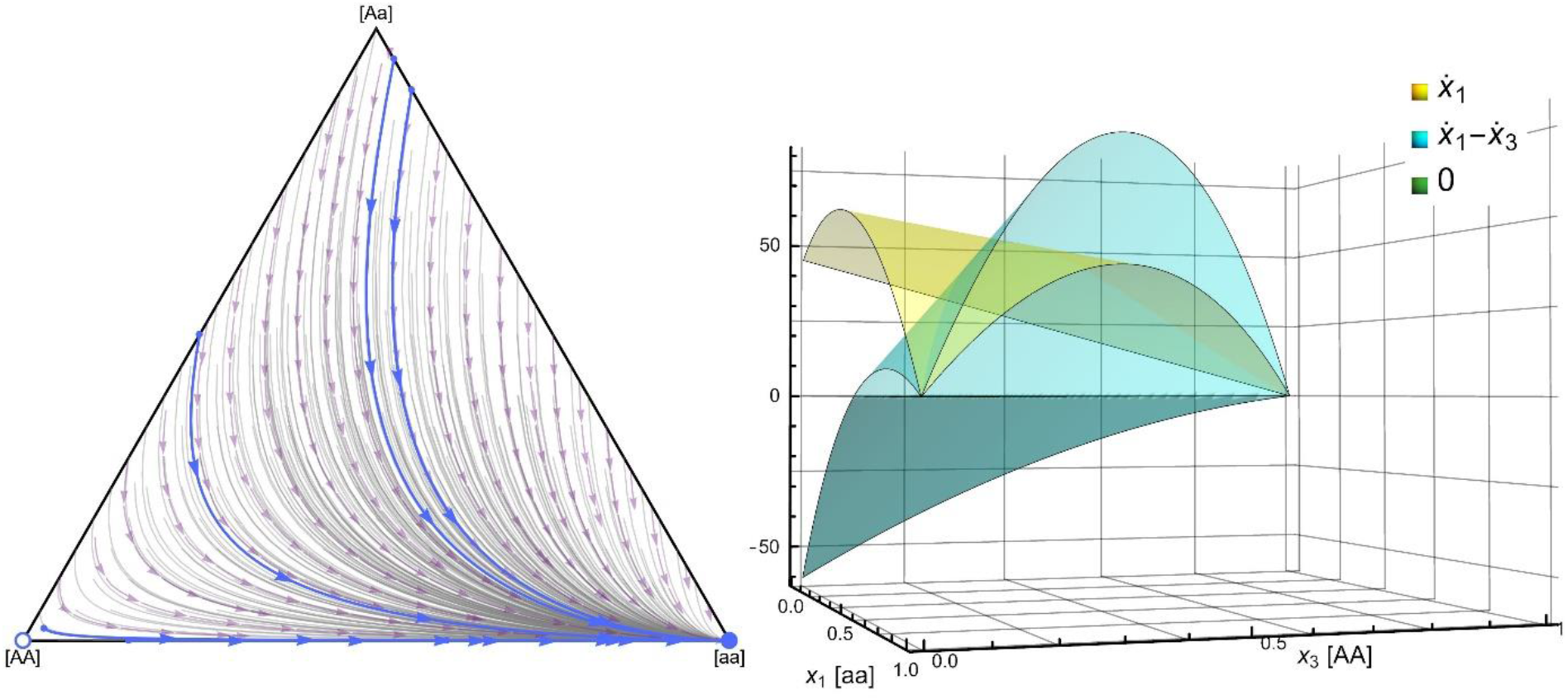
We visualize the genotype dynamics under assortative mating, when the survival probability of the juveniles is determined by a survival donation game. There are *n* = 8 newborns in each family. In Figure 2A, we consider the case where the cooperator allele [a] is recessive, and the basic survival rate *α* = 0.2, the benefit *b* = 0.6, and the cost *c* = 0.1, so the payoff matrix is 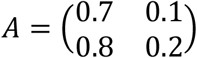. We have already shown that if *b* > *c*, then that cooperator homozygote [aa] is ESGD, therefore locally asymptotically stable. Furthermore, the defector homozygote [AA] is repellor, and this dynamics has no interior rest point, so according to Poincaré–Bendixon theorem, in this selection situation the classical Hamilton rule implies the global stability of the cooperator homozygote [aa].

### Remark 2

Since the equations for 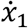 can be rewritten in the following form,

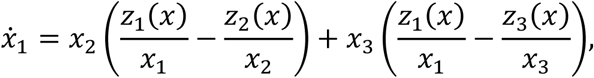

the positivity of 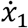 for all *x* ∈ int *S*_3_ can also be written as

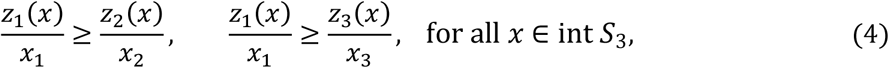

and at least one of these inequalities has to be strict. According to Theorem 2, if inequality (4) is satisfied, the cost-accepting homozygote [aa] is fixed by natural selection. In biological terms, this condition means that *the cost-accepting homozygote is fixed by natural selection provided the other genotypes always receive a smaller average gain from all interactions than the cost-accepting homozygote*.

To see a necessary condition for Remark 2, we show that if all offspring participate in a well-mixed survival game in the entire population before reaching sexual maturity, the cooperator homozygote is never evolutionarily stable in the panmictic population, regardless of whether [a] is dominant or recessive (see SI B). Thus, our conditions (3) and (4) require that the interaction between genotypes is not well-mixed. For example, it is necessary that the number of interactions between two cost-acceptors is greater than the number of interactions between pairs of cost-acceptors and cost-avoiders. Our conditions are consistent with the ideas of Hamilton (1962a,b), who proposed two mechanisms for kin selection. Kin recognition allows individuals to be able to identify their closest relatives carrying identical genes. This idea was further developed by Dawkins (1976): “*I have a green beard and I will be altruistic to anyone else with green beard*.” The essence of Dawkins’s idea is that the person accepting the help is not necessarily the closest relative of the donor, but they share the same genes. Hamilton (1962a,b) also studied viscous populations, where individuals cannot move far from their birthplace, so interactions between relatives are more frequent. The essence of Hamilton’s two mechanisms is that the interaction between individuals is not well-mixed: interactions between relatives are more frequent. Another example of kin selection is Haldane’s familial selection (Haldane 1924), where only the closest relatives, full siblings, interact, and Mendelian inheritance with the mating system determines the phenotypic composition of each family (e.g. Garay et al. 2019, 2023) and the interaction scheme (see Tables 1 and 2).

## 3. Method based on the Price equation

Price (1970) was interested in how the relative frequency of an allele changes from the parent population to the offspring population. We briefly recall his setup and method. He considered the general selection situation, where the production functions of the genotypes can be arbitrary continuous functions on the simplex *S*_3_, see Table 3.

**Table 3.** Selection situation considered by Price.

| Genotype | [aa] | [aA] | [AA] |
| --- | --- | --- | --- |
| Number of allele [a] in genotype | 2 | 1 | 0 |
| Frequency of allele [a] in genotype | 1 | $\frac{1}{2}$ | 0 |
| Frequency of genotype | $x_1$ | $x_2$ | $x_3$ |
| Production | $z_1(x)$ | $z_2(x)$ | $z_3(x)$ |

Note that Price’s selection situation is the same as discussed above. Consequently, the methodological tools of the population genetics method detailed above (ESGD definition, genotype dynamics, etc.) can also be applied. The total production is

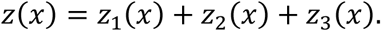

The frequencies of alleles [a] and [A] in the parental population are

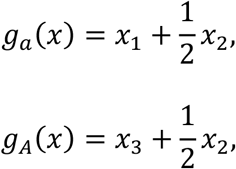

respectively. Clearly, *g*_*a*_(*x*) + *g*_*A*_(*x*) = 1. The genotype distribution in the offspring population is

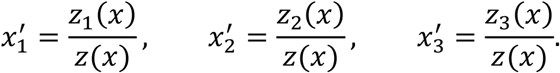

Examining the change in genotype distribution, we obtain the following dynamics:

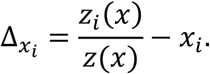

From this dynamic, we derived the genotype dynamics (Garay et al. 2019). To our knowledge, this dynamic has not been studied in the Price equation literature. We believe that this is due to the gene-centric approach of classical population genetics and the gene-centric view of the classical Hamilton rule.

Price was interested in the change of allele frequency. The frequencies of alleles [a] and [A] in the next generation are

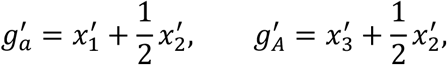

respectively. Now, the well-known Price equation (1970) describes the change in the frequency of allele [a] from a diploid generation to the next generation as follows

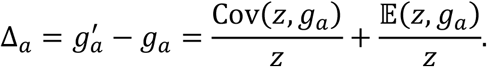

Instead of using the Price equation, we focus on what it is used for (e.g. Frank 1998, van Veelen 2025). Price was interested in how the relative frequency of the allele [a] increases from generation to generation. We can call this *Price monotonicity*: for the allele [a] to become fixed Δ_*a*_ must be strictly positive for all *x* ∈ int *S*_3_. Formally, *Price monotonicity* requires that

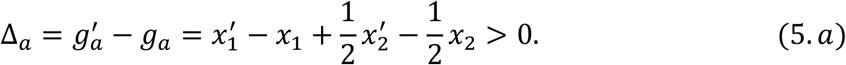

Since Δ_*a*_ + Δ_*A*_ = 0, (5.a) is clearly equivalent to

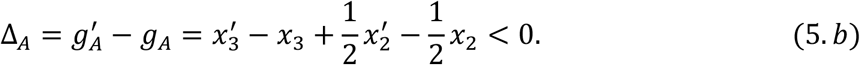

### Remark 3

Price monotonicity is equivalent to Haldane monotonicity. Indeed, Δ_*a*_ > 0 is equivalent to Δ_*a*_ > Δ_*A*_, that is,

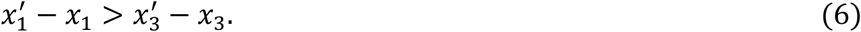

In biological terms, the relative frequency of the allele [a] is increasing if the relative frequency of the homozygote [aa] increases more from generation to generation than that of the homozygote [AA]. Multiplying (6) by the total production *z*(*x*) we arrive at the Haldane monotonicity

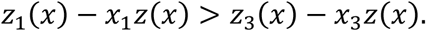

In summary, the condition for monotonicity is the same for the Price equation-based method and the population genetic method, as they impose the same requirement at the diploid stage, which guarantees the global fixation of the cost-accepting homozygote.

### Remark 4

The Price monotonicity does not imply the evolutionarily stability of the cost-accepting homozygote, because it is easy to find *x*_1_, *x*_2_, *x*_3_ and 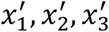 such that 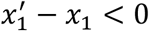, and (4) still holds.

We can now summarize how the two methods relate to each other. Price equation is built on the bases of genotype dynamics. Price monotonicity is equivalent to Haldane monotonicity, so the genotype dynamics can handle all cases that the Price equation does. Moreover, the homozygote [aa] can be globally asymptotically stable in the case where genotype dynamics is not globally Price monotone (see Figures 1 and 2, Theorems 1 and 2).

What is the essential difference between these methods? Price’s method is allele-based, in the sense that the Price equation describes the change in the frequency of the allele [a], and we are interested in the fixation of the allele [a]. Genotype dynamics follows the change in the frequency of the genotype, and we are interested in the stability of its rest points, i.e., the endpoint of natural selection. For example, genotype dynamics can be used to handle both the coexistence of genotypes (see Fig. 1) and bistability, where both homozygote vertices are locally stable (for numerical examples, see Garay et al. 2025).

## 4. Assortative mating

We would like to use an illustrative selection situation to show that when the interaction scheme changes, the conditions for the fixation of cost-accepting homozygotes also change. Here, we only change the mating system of the parents in the selection situation discussed earlier in Example 1. Following Dawkins (1976), we assume that all males with genotypes [aa], [aA], and [AA] have blue, green, and yellow beards, respectively, and that each female accepts only males with her own genotype. In this extremely assortative mating system, the frequency of mating pairs is equal to the frequency of genotypes. The mating table again shows the interaction scheme. Recall that *p*_*k*(*ij*)_ denotes the probability that parents with genotypes *i* and *j* will produce offspring with genotype *k* (see Table 4). For example, if each family has *n* juveniles playing with each other, then the number of the interactions between two genotypes [aa] is 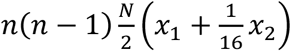 between genotypes [aa] and [AA] is 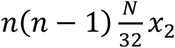, and the number of the interaction between two genotypes [aA] is 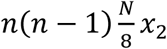, and so on.

**Table 4.** The interaction scheme between juvenile genotypes is determined by the assortative mating system and Mendelian inheritance.

| Genotypes of parents | Average number of couples | genotypes $G_1, G_2, G_3$ | | |
| --- | --- | --- | --- | --- |
| | | $G_1 = [aa]$ | $G_2 = [aA]$ | $G_3 = [AA]$ |
| $G_1 \times G_1$ | $\frac{N}{2}x_1$ | $p_{1(11)} = 1$ | $p_{2(11)} = 0$ | $p_{3(11)} = 0$ |
| $G_2 \times G_2$ | $\frac{N}{2}x_2$ | $p_{1(22)} = \frac{1}{4}$ | $p_{2(22)} = \frac{1}{2}$ | $p_{3(22)} = \frac{1}{4}$ |
| $G_3 \times G_3$ | $\frac{N}{2}x_3$ | $p_{1(33)} = 0$ | $p_{2(33)} = 0$ | $p_{3(33)} = 1$ |

The difference in mating systems has two consequences. The interaction schemes of assortative and random mating are different (see Tables 1 and 4). However, in both mating systems, the probability of interaction between two sibling genotypes depends on the genotype distribution of the parental population; moreover, the interaction between the young is not well-mixed, since now only siblings interact. The mating system also affects the production pattern. For example, in this assortative mating system, two different homozygotes cannot produce heterozygotes because they do not mate. All other parameters of the selection situation are the same as those considered in Example 1. Table 5 presents the selection situation.

**Table 5.** Mating table for a mating system where only individuals with the same genotype form pairs. Each sibling plays exactly one game; the pairs are formed randomly. Here s_1_, s_2_ and s_3_ are the pure strategies of genotypes [aa], [aA] and [AA]. Depending on whether the allele [a] is dominant or recessive, we have s_2_ = s_1_ or s_2_ = s_3_, respectively. Furthermore, n_k(ij)_ is the average number of surviving offspring of genotype k in monogamous families where the parents have genotypes i and j. Finally, A is the payoff matrix.

| Parental genotypes | Average number of couples | Average number of surviving offspring |  |  |
| --- | --- | --- | --- | --- |
| | | $G_1 = [aa]$<br>$s_1$ | $G_2 = [aA]$<br>$s_2$ | $G_3 = [AA]$<br>$s_3$ |
| $G_1 \times G_1$ | $\frac{N}{2}x_1$ | $n_{1(11)} = ns_1As_1$ | $n_{2(11)} = 0$ | $n_{3(11)} = 0$ |
| $G_2 \times G_2$ | $\frac{N}{2}x_2$ | $n_{1(22)} = \frac{n}{16}(s_1As_1 + 2s_1As_2 + s_1As_3)$ | $n_{2(22)} = \frac{n}{8}(s_2As_1 + 2s_2As_2 + s_2As_3)$ | $n_{3(22)} = \frac{n}{16}(s_3As_1 + 2s_3As_2 + s_3As_3)$ |
| $G_3 \times G_3$ | $\frac{N}{2}x_3$ | $n_{1(33)} = 0$ | $n_{2(33)} = 0$ | $n_{3(33)} = ns_3As_3$ |

The production functions are as follows.

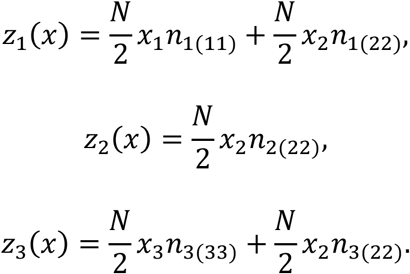

In this selection situation, if *b* > *c* then the homozygote [aa] vertex, the state (1,0,0), is globally stable. To see this, we need the following statements:

a. If *b* > *c*, then the homozygote [aa] is ESGD, regardless of whether allele [a] is dominant or recessive, so the homozygote [aa] vertex is locally asymptotically stable (see SI C2). We have two comments on this statement. First, in the neighborhood of the homozygote [aa] vertex, the genotype dynamics is locally Haldane monotone (see SI C3). Second, this assortative mating system is monogamous, thus only full siblings interact. Therefore, their coefficient of genetic relatedness is *r* = 1/2. Hence, Hamilton’s rule does not apply in this assortative mating. The higher frequency of interactions between homozygotes appears to weaken the indirect effect of genetic relatedness. This may be because if a homozygote [aa] receives help, this help promotes the simultaneous survival of two alleles [a].
b. If *b* > *c*, then the homozygote [AA] vertex is a repellor, regardless of whether allele [a] is dominant or recessive (see SI C4).
c. If *b* > *c*, then there is no interior rest point of the genotype dynamics, regardless of whether allele [a] is dominant or recessive (see SI C5 for the proof). Furthermore, if all genotypes are viable, then the edges of the simplex are repulsive, so there are no rest points on the edges.

Again, Theorem 1 implies the global fixation of the cooperator homozygote. We emphasize that in this selection situation only the inequality *b* > *c* implies the fixation of the cooperator homozygote.

We are also interested in whether Haldane monotonicity holds in the neighborhood of (0,1,0). The answer is negative (see SI C6). This means that we cannot see the global fixation of homozygote [aa] using the Price equation, but we can see it using the genotype dynamics. We emphasize that this system is locally Haldane monotone in the neighborhood of (1,0,0) as well (see Fig. 2).

In Figure 1B, we visualize both monotonicity conditions. This system is not globally Haldane monotone, since 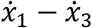 is negative in the neighborhood of the heterozygote vertex (0,1,0). Note that since Price monotonicity and Haldane monotonicity are equivalent, the Price equation cannot show the global stability of homozygote [aa]. However, this case is also locally Haldane monotone in the neighborhood of the heterozygote vertex (0,1,0). Furthermore, our monotonicity conditions hold for this system, since 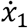 is always positive, so the relative frequency of the homozygote [aa] always increases in the simplex. Thus, we have two independent proofs of the global stability of the homozygote [aa] vertex.

## 5. Discussion

Price equation and genotype dynamics are two methods for studying the fixation of one allele by natural selection in a diploid population. When an allele is fixed, only its homozygotes remain in the diploid population. Using genotype dynamics, we can formulate sufficient conditions for the fixation of cost-accepting behavior. Namely, we give two strict monotonicity conditions for the fixation of a homozygote. The first focuses on the allele distribution, while the second focuses on the genotype distribution.

The genotype dynamics is Haldane monotone if the relative frequency of an allele always strictly increases along all solutions of genotype dynamics, so this allele is fixed (Garay et al 2025). Here we have shown that the genotype dynamics is Haldane monotone if and only if the right-hand side of the Price equation is always strictly positive (see Remark 3). Both conditions guarantee that the relative frequency of an allele is strictly increasing from generation to generation. This equivalence also means that any result that can be obtained using the Price equation can also be obtained using the genotype dynamics. The converse is not true. This is because Haldane monotonicity is a rather strict global condition for the global stability of a homozygote (see Figures 1 and 2). Note that in the neighborhood of the homozygote vertex, the genotype dynamics is locally Haldane monotone (see Figures 1 and 2).

Our monotonicity condition (see Theorem 2 and Remarks 1, 2) needs that the relative frequency of a homozygote always strictly increases according to the genotype dynamics, e.g. the cost-accepting homozygote is fixed by natural selection if the other genotypes always receive a smaller average gain from all interactions than the cost-accepting homozygote. We have found that these two monotonicity conditions are not equivalent (see Remark 1).

Our monotonicity condition requires that the interaction is not well-mixed in the diploid population. Hamilton (1962a,b) has already proposed two mechanisms for kin selection: kin recognition and viscous populations. In familial selection, the interaction is also not well-mixed (see Tables 3 and 6), and our example belongs to this selection situation. It is not surprising that even in a diploid population, a necessary condition for the survival of the cost-accepting type is that the interaction is not well-mixed. We note that the interaction scheme has an elementary effect in asexual classical evolutionary game theory, as well. Consider, for example, an asexual population and the individuals play a game. If the probability of interaction within the same phenotype is high enough, then the verbal definition of evolutionary stability implies that the phenotype maximizes the average payoff of the interaction (see Section 5 in Garay & Varga 2011). In other words, if the interaction scheme is far from the well-mixed interaction, then the game-theoretic method cannot be applied in evolutionary theory. We also showed that if the interaction is well-mixed in the diploid population and the juveniles play a survival game, then the cooperator homozygote is not fixed.

Furthermore, we have given non-monotonicity conditions that imply the fixation of a homozygote (see Theorem 1). The fixation of a homozygote depends on the phenotype-dependent payoff of the interaction (Examples 1 and 2, see also Garay et al. 2025), the genotype-phenotype mapping (Garay et al. 2025), the interaction scheme (see Tables 1, 4) and the mating system (Examples 1 and 2, see also Garay et al. 2023). We give examples to show that changing only the mating systems, namely, if we consider panmixing or assortative mating systems, the condition for the fixation of the cooperator homozygote [aa] also changes: from *b* > 2*c* (see Example 1) to *b* > *c* (see Example 2)

In the Introduction, we have already mentioned that there is a controversial opinion that one of the drawbacks of the Price equation is its dynamic insufficiency (e.g. Frank 1995, van Veelen et al. 2012). Here we find that the Price equation is not dynamically insufficient from the point of view of genotype dynamics, since Price monotonicity is equivalent to Haldane monotonicity of genotype dynamics. This means that Price’s method gives the same sufficient conditions as Haldane monotonicity, but these are stronger than necessary conditions for homozygote fixation. In fact, our non-monotonicity conditions (Theorem 1) and our monotonicity conditions (Remarks 2, 3) also imply homozygote fixation.

## Acknowledgments

We are grateful to István Zachár for his comments and the figures and to Tamás Varga for his comments.

## Supporting Information

### SI.A. Application of the Poincaré–Bendixson theorem

Consider the distribution of genotypes belongs to the 2-dimensional simplex

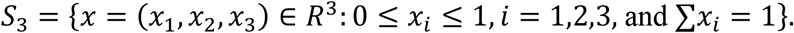

Denote the relative frequency of genotype [aa], [aA] and [AA] is *x*_1_, *x*_2_ and *x*_3_ in the parent population, respectively. Let us define the *production* functions *z*_1_, *z*_2_, *z*_3_ as the number of genotypes [aa], [aA] and [AA] in the next parent population; (*z*_1_, *z*_2_, *z*_3_): *S*_3_ → *R*^3^. We can assume they are smooth. Let us consider the genotype dynamics

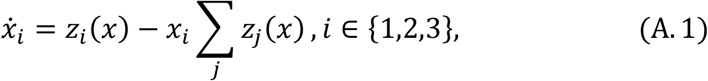

### A.1.

#### Theorem 1

Application of the Poincaré–Bendixson theorem. Assume that

(d) the homozygote [aa] vertex is locally asymptotically stable,
(e) the homozygote [AA] vertex is a repellor,
(f) Besides the homozygous verteces (1,0,0) and (0,0,1), respectively, the genotype dynamics (A.1) has no rest point.

Then (1,0,0), that is the homozygous [aa] state, is a globally asymptotically stable rest point of the genotype dynamics.

*Proof*. To see this, apply the Poincaré–Bendixson theorem according to which a bounded solution converges to either an equilibrium point, a periodic solution, or a so-called graphic (e.g., a heteroclinic cycle, a homoclinic cycle, or a union of such trajectories connected at equilibrium points) (Theorem 4.5.1 in Kong [2] or Ch. 3.7 in Perko [3]). Since a periodic orbit must contain an equilibrium point in its interior (Theorem 4.7.2 in Kong [2] or Corollary 2 in Ch. 3.12 of Perko [3]), a periodic solution cannot exist, because we assumed that no equilibrium points exist other than (0,0,1) and (1,0,0).

Since (1,0,0) is asymptotically stable, while no solution tends to (0,0,1), neither a homoclinic cycle nor a heteroclinic cycle can exist (the latter would require a solution that starts from (1,0,0), which is impossible since (1,0,0) is stable). Consequently, no graphic can exist on the simplex.

It follows that solutions can tend to only one of the two equilibrium points. Since, by assumption, no solution tends to (0,0,1), every solution other than the constant solution (*x*(*t*), *y*(*t*), *z*(*t*)) = (0,0,1) must tend to (1,0,0) as *t* tends to +∞. This is precisely the statement that (1,0,0) is globally stable.

### A.2.

#### Theorem 2

Monotonicity condition on genotypes. Suppose that 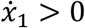 on the set *S*_3_\{(1,0,0), (0,0,1)}. Then (1,0,0) is asymptotic stable.

*Proof* Condition 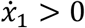 implies that *L*(*x*) = *x*_1_ is a Lyapunov function at (1,0,0) on the simplex *S*_3_, since it reaches its maximum at the point (1,0,0) on *S*_3_ and its derivative

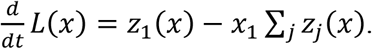

along the differential equation (A.1) is positive. This implies the asymptotic stability of (1,0,0) (p.194 in Hirsch et al. [1] or Theorem 3.5.1(b) in Kong [2]). Geometrically, this case means that a trajectory can intersect any line parallel to the edge between (0,0,1) and (0,1,0) at most once, and that no such line can be tangent to a trajectory.

Note that if 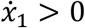 holds for every *x* ∈ *S*_3_\{(1,0,0), (0,0,1)}, then the previous argument with the Lyapunov function works on the entire simplex, which implies global asymptotic stability of (1,0,0) without using Theorem 1.

### SI.B. Well mixed interaction between genotypes

The offspring participate in a survival game before reaching sexual maturity. Let the interaction be well-mixed, that is, interaction is not only possible within the family.

#### B.1. Basic quantities

Let *h*_*i*_(*x*) denote the proportion of genotype *i* newborns in the parental population being at state *x*. The total number of newborns is denoted by *H*(*x*). The offspring participate in a survival game before reaching sexual maturity. Let the interaction be well-mixed. Then each adolescent individual has probability *h*_*i*_(*x*) of interacting with an individual of genotype *i*.

Assume panmixia. The proportion of different genotypes among the newborns can be computed from the following mating table.

**Table 1.**
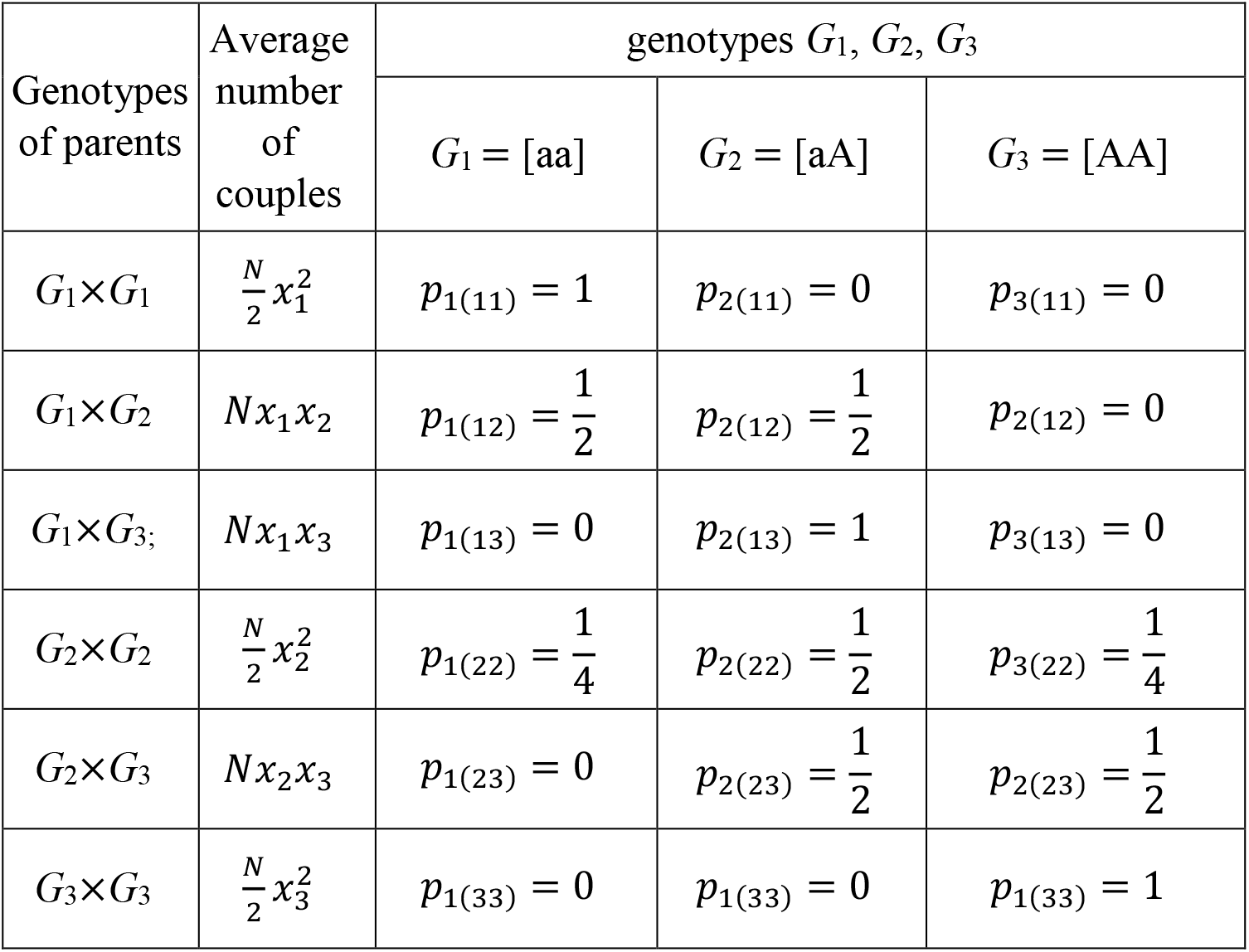
Mendelian inheritance determine the phenotypic composition of each family.

Let *B*_*i*_(*x*) denote the average benefit that an individual of type *i* receives in a population at state *x*. Similarly, *C*_*i*_(*x*) denote the average cost it has to pay.

There are two cases according that [a] is dominant or recessive.

In the **dominant** case [aA] is also cost-accepter, thus

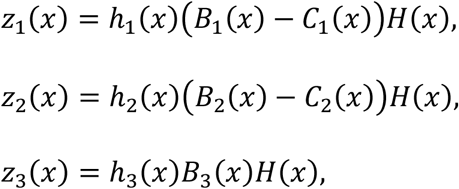

Recall that multiplying the right-hand sides by **the same** positive function of *x* is equivalent to a time transformation, i.e., we switch from *x*(*t*) to *x*(*φ*(*t*)) with a suitable increasing function *φ*. Such a transformation does not affect the asymptotic behavior of the dynamical system. Thus the multiplier *H*(*x*) can be ignored, but we will keep it in the following to avoid misunderstandings.

Since each individual has probability 1 − *h*_3_(*x*) to interact with a cooperator, we have

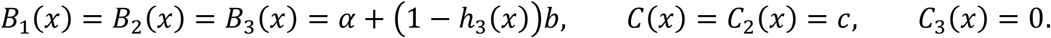

Let *B*(*x*) denote the common value of *B*_1_(*x*) = *B*_2_(*x*) = *B*_3_(*x*). Then

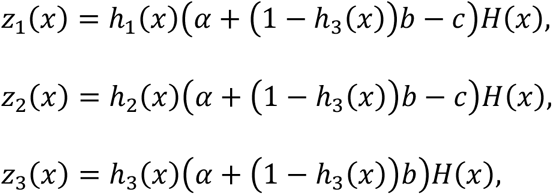

therefore

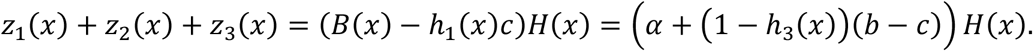

This is the average portion of survivors in the newborn population: the base survival probability modified by the benefit-cost balance of the cost-accepter individuals.

If [a] is **recessive**, then [aA] is cost-avoider. This time each individual has probability *h*_1_(*x*) to interact with cost-accepter homozygote, thus

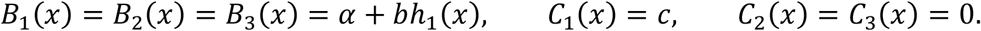

Furthermore,

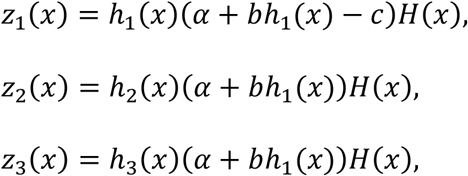

and

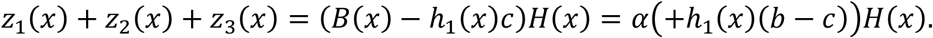

Assume that the same number of offspring is born in each family, say *n*. Then 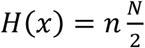, and panmixia implies

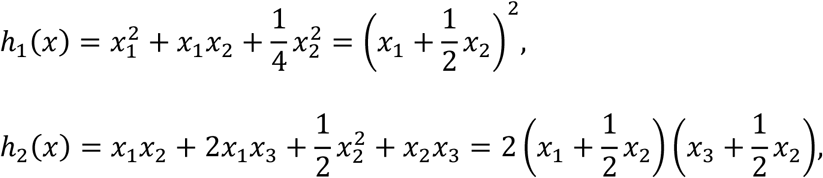

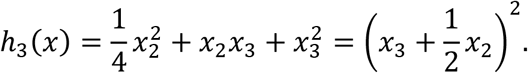

#### B.2. Homozygote [aa] is not evolutionarily stable

There are two cases according that [a] is dominant or recessive.

In the **recessive** case we have

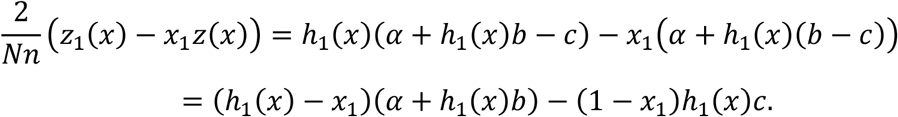

We will show that 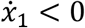 in a sufficiently small neighborhood of the state (1,0,0).

Let 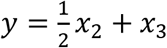, fixed and sufficiently small. Clearly, *x*_3_ can be arbitrary in the interval (0, *y*). Now, *h*_1_(*x*) = (1 − *y*)^2^, *h*_1_(*x*) − *x*_1_ = *y*^2^ − *x*_3_, and 1 − *x*_1_ = 2*y* − *x*_3_. Thus

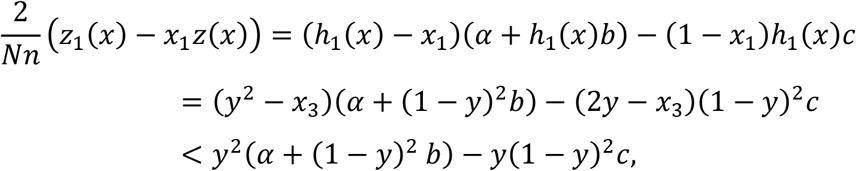

which is negative if *y* is small enough. Thus the state (1,0,0) is a repellor.

Summary: In the recessive case 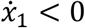 in a sufficiently small neighborhood of the state (1,0,0). Thus the state is a repellor.

Let us turn to the **dominant** case. We will show that in every environment of (1,0,0) there is a state where 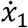 is negative and a state where 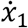 is positive. This time

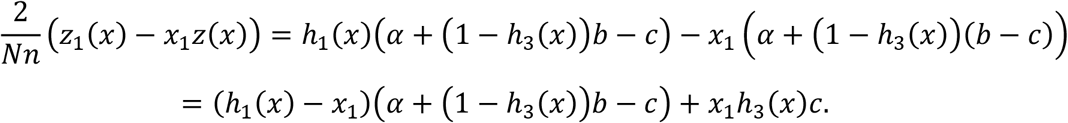

With the notations of the recessive case we have *h*_3_(*x*) = *y*^2^, hence

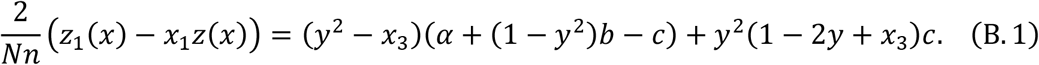

Let *x*_3_ tend to zero first. Then the right-hand side of (B.1) converges to

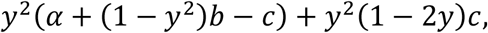

which is positive. On the other hand, if *x*_3_ → *y*, then the right-hand side of (B.1) converges to

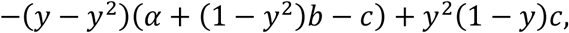

which is negative, provided *y* is small enough.

Summary: In the dominant case, in every environment of (1,0,0) there is a state where 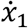 is negative and a state where 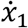 is positive.

### SI.C. Assortative mating

#### C.1. Basic quantities

Mating pairs consist exclusively of individuals with the same genotype. We suppose that there are *n* newborns in each family. Let *n*_*i*(*jj*)_ denote the average number of genotype *i* offspring in a family of genotype *j*. Genotypes [aa], [aA], [AA] are numbered 1, 2, 3, respectively. Suppose the benefit and the cost are 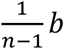 and 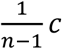 in every interaction where a cost accepter participates. Clearly,

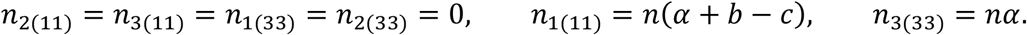

Firstly, let us deal with the case where the cost accepter behavior is **dominant**.

Consider a heterozygote family. Let *ξ*_1_, *ξ*_2_, *ξ*_3_ denote the random number of newborns with genotypes 1, 2, and 3, respectively. The joint distribution of (*ξ*_1_, *ξ*_2_, *ξ*_3_) is multinomial 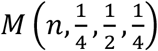. Then we have

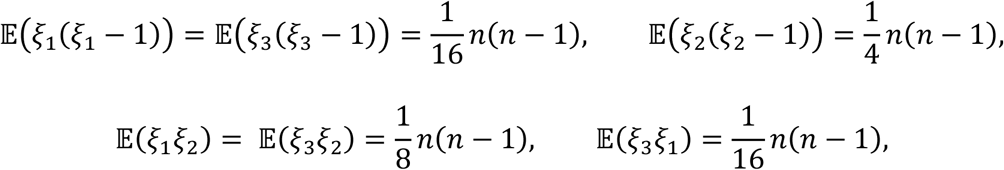

where 𝔼 stands for expectation. Hence

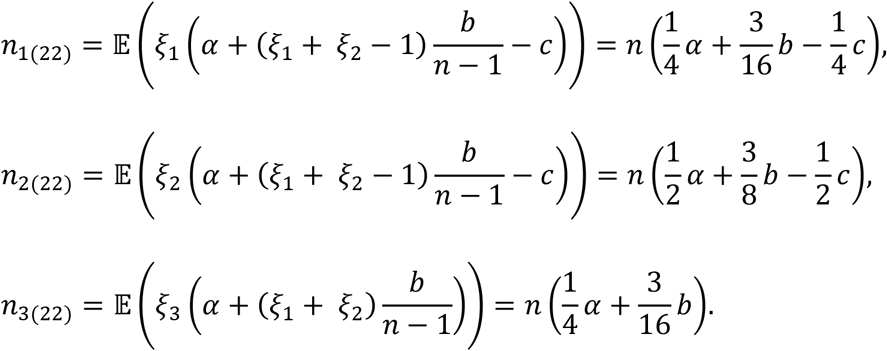

The average production of different genotypes in the next generation can be computed by using the formula 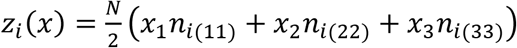. Thus,

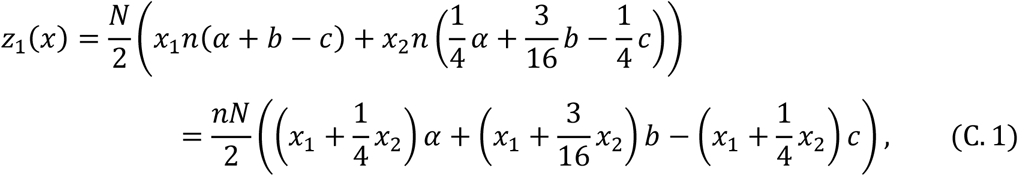

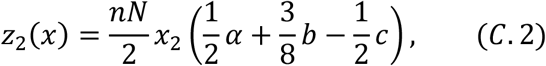

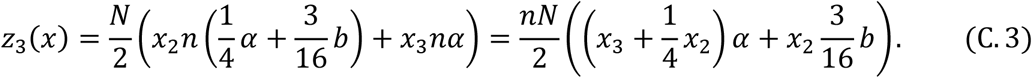

The total production, i.e., the size of the next generation is

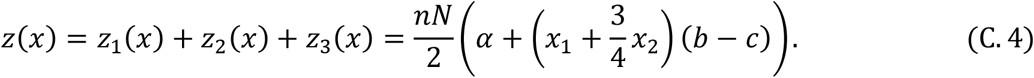

Let us calculate the change in the frequency of each genotype, projected to the size of the offspring generation, i.e., the quantities *z*_*i*_(*x*) − *x*_*i*_*z*(*x*). From (C.1)–(C.4) we get

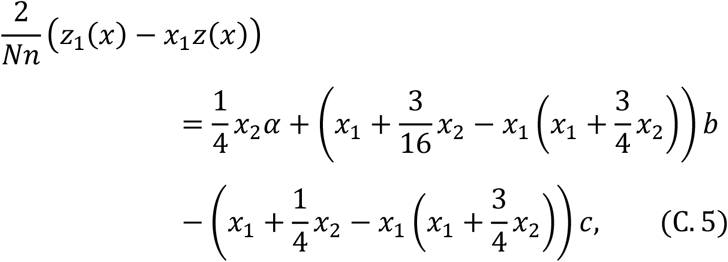

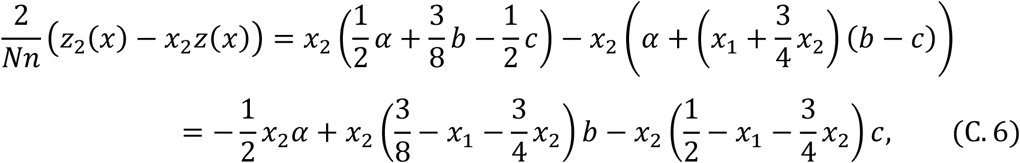

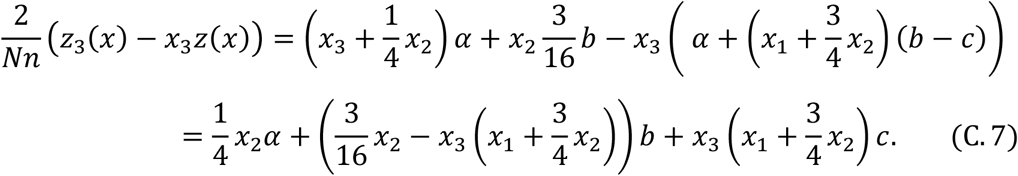

Next, let us turn to the case where the cost accepter behavior is **recessive**. Then nothing will change but *n*_*i*(22)_, *i* = 1, 2, 3. This time

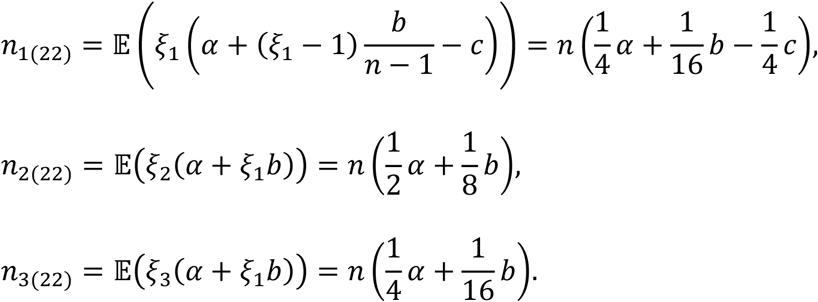

Hence, the average production of different genotypes in the next generation is

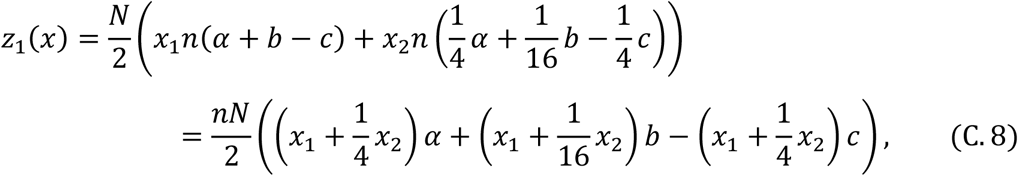

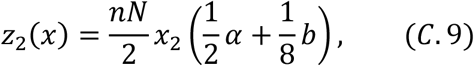

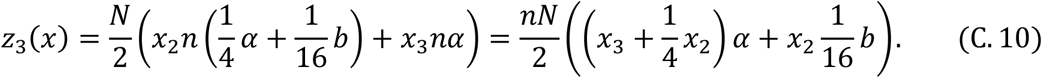

The total production is

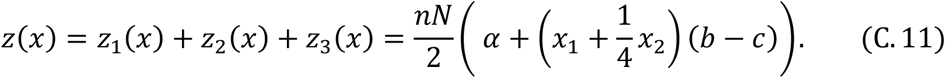

From (C.8)–(C.11) we obtain

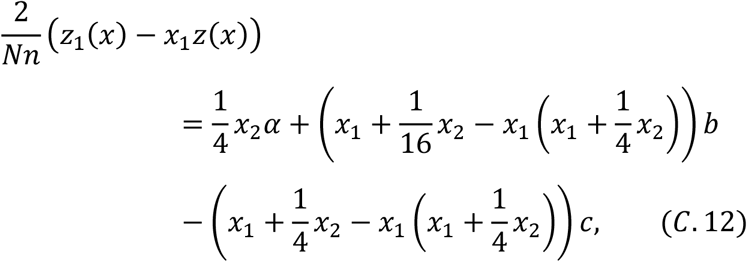

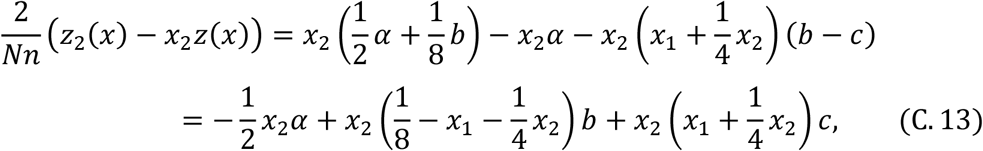

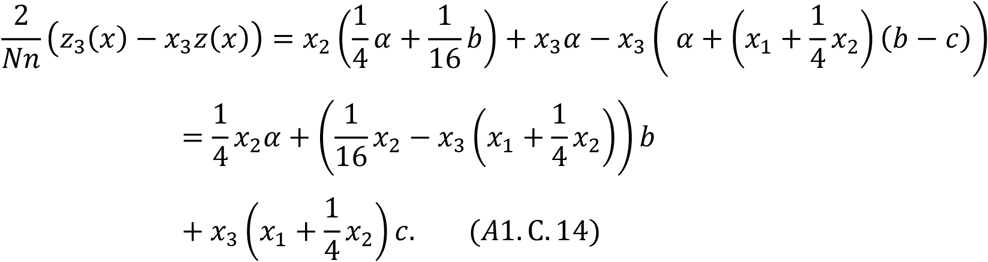

#### C.2. Condition for the homozygote [aa] to be evolutionarily stable

Genotype [aa], or in other words, state *x*^*^ = (1,0,0) is an ESDG, if *z*_1_(*x*) > *x*_1_*z*(*x*), whenever *x* is sufficiently close to *x*^*^.

**Dominant** case. Let us start from (C.5).

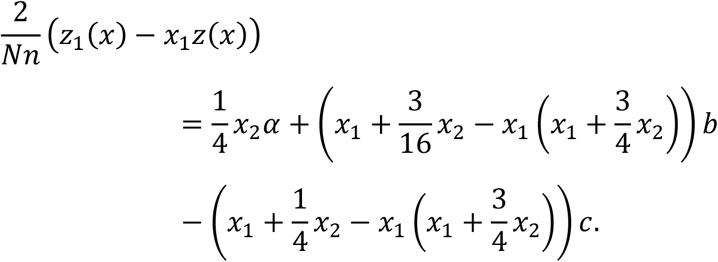

Here

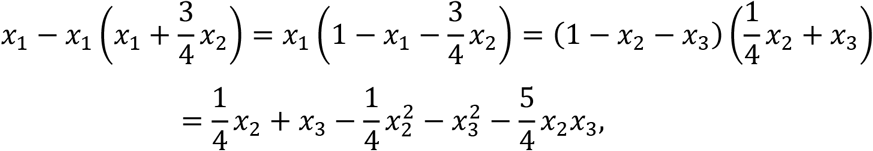

therefore

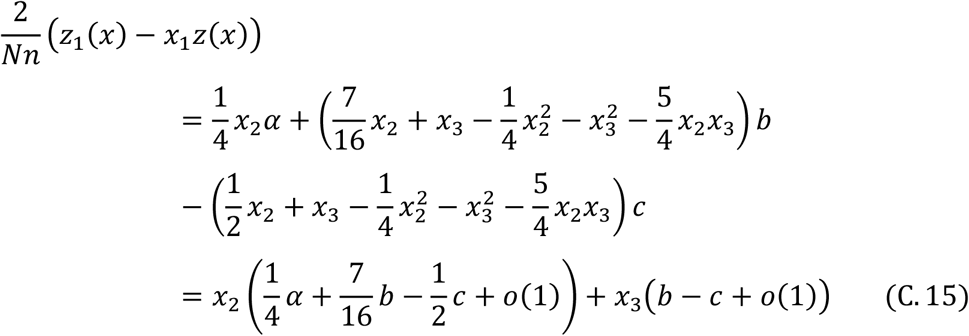

as *x*_2_ and *x*_3_ tend to zero. Here *o*(1) denotes a quantity that tends to zero as *x*_2_ and *x*_3_ do so. Since the ratio of *x*_2_ and *x*_3_ can be any, for the positivity of *z*_1_(*x*) − *x*_1_*z*(*x*) it is necessary that the coefficients of *x*_2_ and *x*_3_ are also nonnegative (and their positivity is already sufficient). That is, conditions *b* > *c*, and 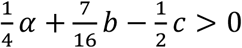 will do. In order that our formula for the survival probability remain between 0 and 1 we need that 0 ≤ *α* − *c* and *α* + *b* ≤ 1. Typically *b* and *c* are small and the basic survival probability *α* is not close to 0 or 1, therefore this constraints are not unrealistic. In light of this,

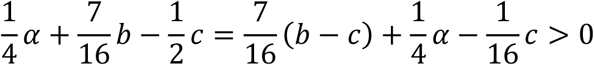

holds whenever *b* > *c*. Thus *x*^*^ = (1,0,0) is an ESDG, if *b* > *c*.

Next, let us turn to the case where the cost accepter behavior is **recessive**. This time we start from (C.12)

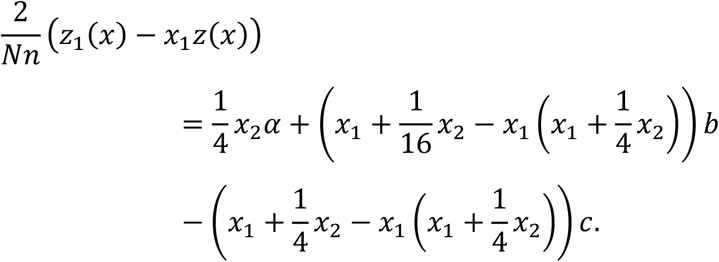

Here

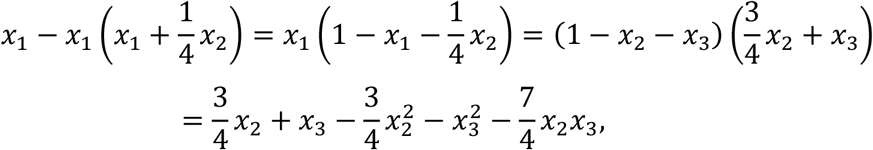

consequently

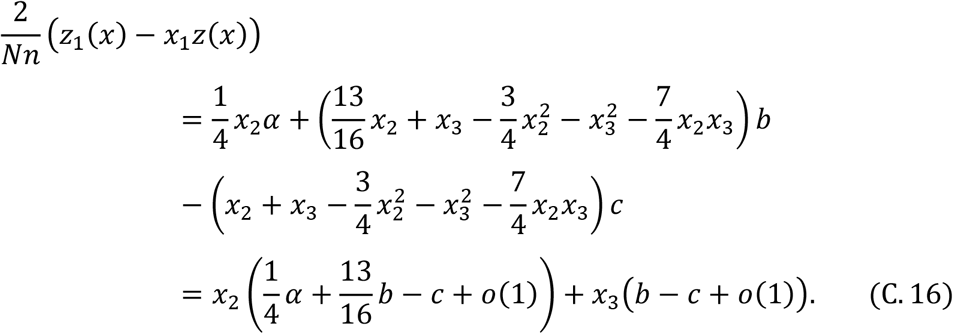

Because of the second term we need *b* > *c* again; in addition, we also need that

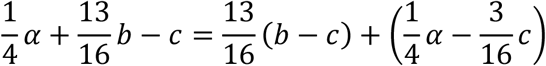

be positive. Since both summands in the right-hand side are positive, this does not impose further conditions. Thus (1, 0, 0) is an ESDG if *b* > *c*.

#### C.3. Conditions for Haldane monotonicity in the neighborhood of (1,0,0)

Case of **dominant** inheritance.

From equation (C.7) it follows that

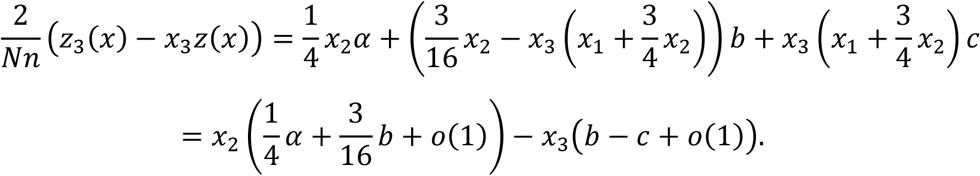

Combining this with (C.15) we get

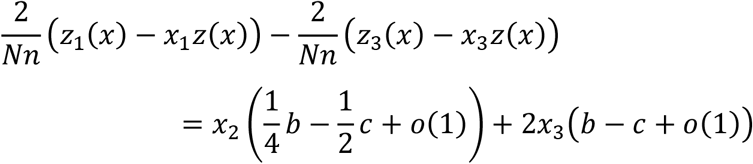

The condition that both the coefficients of *x*_2_ and *x*_3_ are positive as *x*_1_ is close to 1 is *b* > 2*c*.

Case of **recessive** inheritance. Equation (C.14) implies

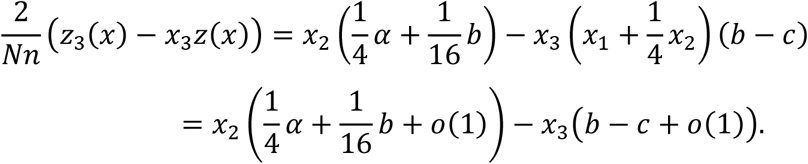

Recalling (C.16) we can write

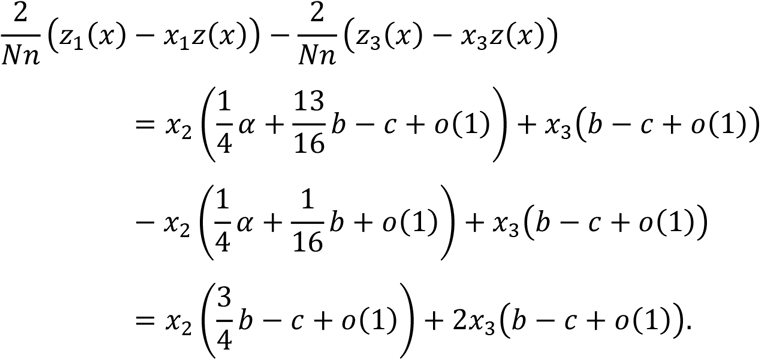

Both the coefficients of *x*_2_ and *x*_3_ are positive as *x*_1_ is close to 1 if 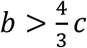.

#### C.4. The homozygote [AA] vertex (state (0,0,1)) is not locally stable

We will show that in a sufficiently small neighborhood of (0,0,1) the frequency of homozygote [aa] strictly increases, regardless of whether allele [a] is dominant or recessive, provided *b* > *c*. Formally, *z*_1_(*x*) − *x*_1_*z*(*x*) > 0, where *z*(*x*) = ∑_*j*_ *z*_*j*_(*x*).

In the **dominant** case, according to (C.5) we have

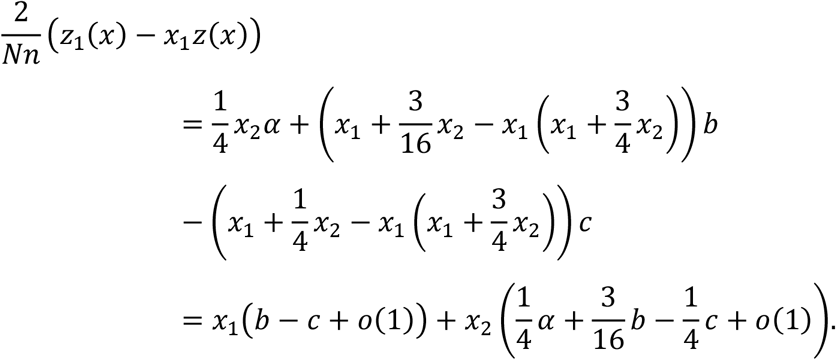

Here *o*(1) denotes a quantity tending to zero as *x*_1_ and *x*_2_ do so. Clearly, both the coefficients of *x*_1_ and *x*_2_ are positive as *x*_3_ is sufficiently close to 1, because *b* − *c* and

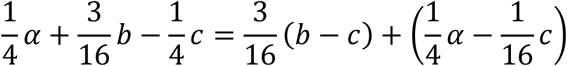

are positive by supposition.

The **recessive** case is quite similar. This time we start from (C.12).

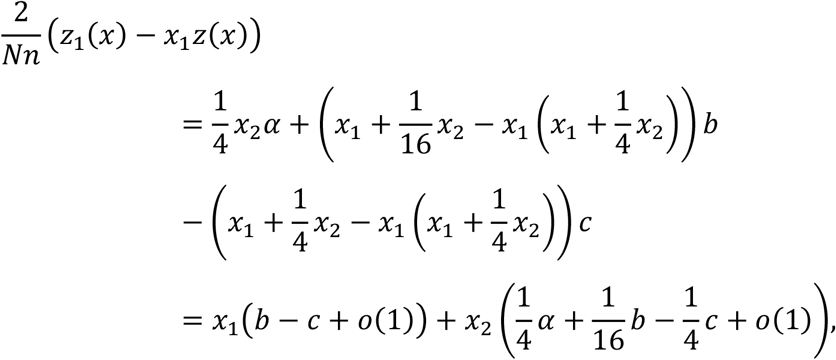

and here both *b* − *c* and

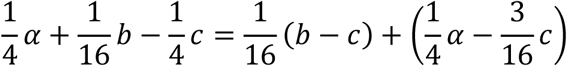

are positive.

#### C.5. There is no interior rest point of the genotype dynamics

If there were an interior rest point, it would satisfy the following equation (Garay et al 2025)

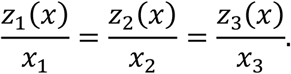

In the **dominant** case, by equations (C.1) and (C.2),

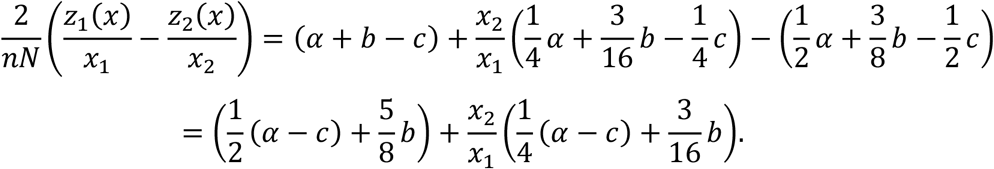

Here both summands are positive, since *α* ≥ *c* by supposition.

In the **recessive** case, by (C.8) and (C.9) we have

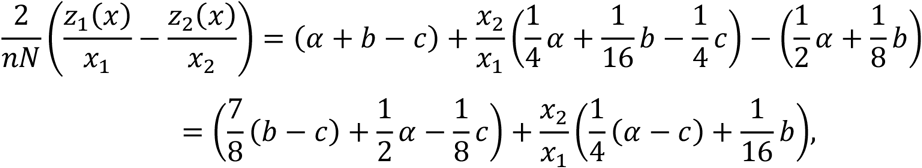

where both summands are positive if *b* > *c*.

#### C.6. Haldane monotonicity does not hold in the neighborhood of (0,1,0)

In the **dominant** case, from (C.5) and (C.7) we have

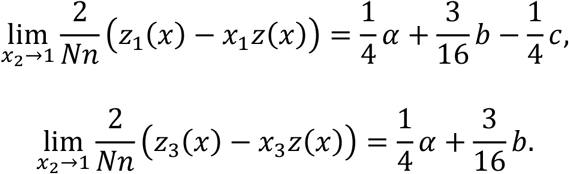

Note that *x*_2_ → 1 implies *x*_1_ → 0 and *x*_3_ → 0. Thus,

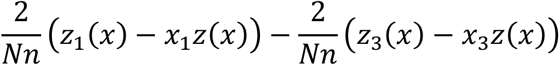

is negative in a sufficiently small neighborhood of (0,1,0).

The **recessive** case is similar. From (C.8) and (C.10) it follows that

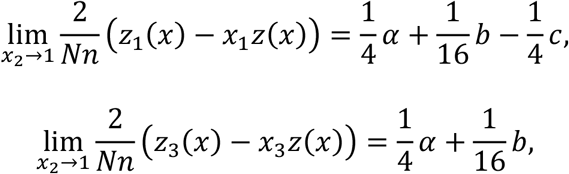

hence

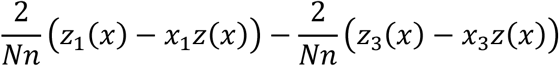

is negative in a sufficiently small neighborhood of (0,1,0).

## Notes

**Competing Interest Statement:** The authors declare that there are no conflicts of interest regarding the publication of this paper.

### Competing Interest Statement

The authors have declared no competing interest.

## REFERENCES

Allen B., Khwaja AR., Donahue JL., Kelly TJ., Hyacinthe SR., Proulx J., Lattanzio Cassidy J., Dementieva A., Sample C. (2024) Nonlinear social evolution and the emergence of collective action, PNAS Nexus 3,3 131, 10.1093/pnasnexus/pgae131

Alós-Ferrer C., Hofbauer J. (2022) Excess payoff dynamics in games. J Econ Theory.; 204: 105464. doi: 10.1016/j.jet.2022.105464

Dawkins R. (1976) The Selfish Gene. Oxford: Oxford University Press. ISBN 978-0-19-217773-5.

Frank SA. (1995) George Price’s contributions to evolutionary genetics. J. Theor. Biol. 175, 373–388 10.1006/jtbi.1995.0148

Frank SA. (1998) Foundations of Social Evolution. Princeton University Press, Princeton

Garay J., Garay BM., Varga Z., Csiszár V., Móri, TF. (2019) To save or not to save your family member’s life? Evolutionary stability of self-sacrificing life history strategy in monogamous sexual populations. BMC Evolutionary Biology 19, 147.

Garay J., Csiszár V., Móri TF. (2023) Subsistence of sib altruism in different mating systems and Haldane’s arithmetic, J. Theor. Biol. 557, 111330, 10.1016/j.jtbi.2022.111330

Garay J., López, I., Varga, Z. Csiszár V., Móri TF. (2024) Survival cost sharing among altruistic full siblings in Mendelian population. BMC Ecol Evo 24, 142 10.1186/s12862-024-02317-z

Garay J.,Varga Z. (2011) Survivor’s dilemma: Defend the group or flee? Theor. Pop. Biol. 80:217–225. 10.1016/j.tpb.2011.08.003

Garay J, Varga T, Csiszár V, Móri TF, Szilágyi A. (2025) Matrix games between full siblings in Mendelian populations. PLoS One 20(9): e0331044. 10.1371/journal.pone.0331044

Gardner A., West SA, Wild G. (2011) The genetical theory of kin selection. J Evol Biol, 24, 1020–1043, doi/10.1111/j.1420-9101.2011.02236.x

Haldane JBS. (1924) A mathematical theory of natural and artificial selection. Part. I. Trans. Camb. Phil. Soc. 23:19–41.

Haldane J.B.S., (1955) Population genetics. New Biology, 18:34–51.

Haldane JBS, Jayakar SD. (1965) Selection for a single pair of allelomorphs with complete replacement. J Genetics. 59:81–7. ISBN9781315768694

Hamilton WD. (1964a) The genetical evolution of social behaviour. I. J. Theor. Biol.;7:1–16.

Hamilton WD. (1964b) The genetical evolution of social behaviour. II. J. Theor. Biol.;7:17– 52.

Hamilton WD. (1970) Selfish and Spiteful Behaviour in an Evolutionary Model. Nature 228, 1218–1220. 10.1038/2281218a0

Hofbauer J, Sigmund K. (1998) Evolutionary games and population dynamics. Cambridge: Cambridge University Press; ISBN-13: 978-0521625708.

Hull P. (1964) Partial incompatibility not affecting total litter size in the mouse. Genetics. 50:563–70. doi: 10.1093/genetics/50.4.563

Maynard Smith, J. (1964) Group Selection and Kin Selection. Nature. 201 (4924): 1145–1147. doi:10.1038/2011145a0

Maynard Smith J, Price GR. (1973) The logic of animal conflict. Nature. 246: 15–18. doi:10.1038/246015a0.

Queller DC. (1985) Kinship, reciprocity and synergism in the evolution of social behaviour. Nature 318:366–367. DOI: 10.1038/318366a0

Queller DC. (1992) A general model for kin selection. Evolution; International Journal of Organic Evolution 46:376–380. DOI: 10.1111/j.1558-5646.1992.tb02045.x, PMID: 28564031

Van Veelen M. (2009) Group selection, kin selection, altruism and cooperation: When inclusive fitness is right and when it can be wrong. J. Theor. Biol. 259:589–600

Van Veelen M. (2025) The general version of Hamilton’s rule. eLife 14:RP105065. 10.7554/eLife.105065.3

Van Veelen M, Allen B, Hoffman M, Simon B, Veller C. (2017) Hamilton’s rule. J Theor Biol. 414: 176–230. doi: 10.1016/j.jtbi.2016.08.019.

Van Veelen M, García J, Sabelis MW, Egas M. (2012) Group selection and inclusive fitness are not equivalent; the Price equation vs. models and statistics. J Theor Biol. 299: 64–80. doi:10.1016/j.jtbi.2011.07.025.

Van Veelen M. (2005) On the use of the Price equation. J Theor Biol. 237: 412–426. doi:10.1016/j.jtbi.2005.04.026.

## References

[1] Hirsch MW, Smale S, Devaney RL. Differential Equations, Dynamical Systems, and an Introduction to Chaos. 2nd ed. San Diego: Academic Press, 2004. ISBN: 9780123497031

[2] Kong Q. A Short Course in Ordinary Differential Equations. Springer, 2014. ISBN: 9783319112398

[3] Perko L. Differential equations and dynamical systems, 3^rd^ ed. Springer, 2001. ISBN: 9780387951164

